# Accurate prediction of protein tertiary structural changes induced by single-site mutations with equivariant graph neural networks

**DOI:** 10.1101/2023.10.03.560758

**Authors:** Sajid Mahmud, Alex Morehead, Jianlin Cheng

## Abstract

Predicting the change of protein tertiary structure caused by singlesite mutations is important for studying protein structure, function, and interaction. Even though computational protein structure prediction methods such as AlphaFold can predict the overall tertiary structures of most proteins rather accurately, they are not sensitive enough to accurately predict the structural changes induced by single-site amino acid mutations on proteins. Specialized mutation prediction methods mostly focus on predicting the overall stability or function changes caused by mutations without attempting to predict the exact mutation-induced structural changes, limiting their use in protein mutation study. In this work, we develop the first deep learning method based on equivariant graph neural networks (EGNN) to directly predict the tertiary structural changes caused by single-site mutations and the tertiary structure of any protein mutant from the structure of its wild-type counterpart. The results show that it performs substantially better in predicting the tertiary structures of protein mutants than the widely used protein structure prediction method AlphaFold.

## 1 Introduction

Genes and proteins in every species undergo constant mutations, which are the driving force of evolution. A single amino acid mutation (i.e., the replacement of one residue with another one) in a protein can change its structure, leading to gain, loss, or adjustment of its function and interaction. Therefore, studying the effects of single-site mutations is important for elucidating the relationships between genotypes and phenotypes. For instance, many single-site mutations are implicated in human diseases such as cancers [1–4]. Thus, it is important to determine the structural changes in proteins induced by single-site mutations.

Experimental techniques such as x-ray crystallography can be used to determine the tertiary structures of protein mutants with single-side mutations, but they can be only applied to a small portion of proteins due to their low throughput and high cost. The cutting-edge protein structure prediction methods such as AlphaFold [5] that revolutionalized structural biology recently can rather accurately predict the overall tertiary structures of most proteins, but they are not sensitive enough to accurately predict the smaller structural change caused by one amino acid change in a protein sequence. That is, given the sequence of a wild-type protein or its mutant with a single amino acid substitution, AlphaFold still cannot accurately predict their structural difference [6, 7]. Therefore, there is a significant need to develop specialized methods to predict the structural changes caused by single-site mutations.

Existing specialized protein mutation prediction methods [8–17] focus on predicting the overall stability change (i.e., a change of Gibbs free energy difference [8, 12]) of protein structure or the functional (e.g., pathogenic) changes caused by mutations [4]), without directly predicting the exact mutation-induced tertiary structural change, partially because the accurate prediction of the tertiary structure of protein mutants has been a much more challenging problem to tackle. Therefore, the prediction of the structure of protein mutants is an outstanding problem that has not been well explored. In this work, we propose to develop a deep learning method based on rotation- and translation-equivariant graph neural networks (EGNN) to directly predict the tertiary structure of protein mutants with a single-site mutation from the structures of their wild-type counterparts.

Deep learning has been widely applied to a variety of problems of predicting protein properties from sequences, such as tertiary structure prediction [5], inter-protein contact and distance prediction [18–20], protein domain boundary prediction [21–23], and protein-ligand interaction prediction [24–28]. A recent trend is to develop deep learning methods to directly take three-dimensional (3D) protein structures as input to predict their properties (e.g., protein function) [29] or further refine them [30]. In this work, we formulate the problem of predicting the structural change induced by a single-site mutation as a problem of directly predicting the 3D structure of a protein mutant from the known 3D structure of its corresponding wild-type protein, capturing the structural change from the wild-type protein to the mutant. Directly using 3D protein structures as input for deep learning is more challenging than using sequence information as input because the coordinates of atoms of the protein structures change when they are rotated and translated, while the properties of the proteins (e.g. structural changes induced by mutations) do not. Therefore, deep learning methods that can capture the essential features of protein structures in the feature space regardless of the rotation and translation are needed to effectively to predict protein properties from protein structure input.

One type of deep graph neural networks (GNNs) [31], known as Equivariant Graph Neural Networks (EGNN) [32], has been recently developed to capture the essential features of 3D objects that are equivariant with respect to rotation, translation, and permutation. An EGNN uses a graph to represent a 3D object, where nodes denote the points of the 3D object and edges denote the distance relationship between the points. Each node has the invariant features not changing with respect to the position of the point (e.g., atom type) and the positional features (e.g., x, y, z coordinate of the point). If the graph of a 3D object is rotated and translated in the input space, the same hidden features can be generated by the EGNN in the equivariantly rotated and translated position in the hidden feature space.

Based on the EGNN and residual network [33], we developed a method called PreMut to predict the coordinates of all the atoms of a protein mutant with a single-site mutation from the known tertiary structure of its corresponding wild-type protein. To the best of our knowledge, PreMut is the first deep learning method designed to directly predict tertiary structural changes caused by single-site mutations and the tertiary structure of protein mutants. To train and test PreMut, we curated several protein mutant structure datasets from the Protein Data Bank (PDB) [34]. The benchmark results show that PreMut can rather accurately predict the tertiary structures of protein mutants and performs better than the state-of-the-art protein structure prediction method AlphaFold.

## 2 Results

### 2.1 The Equivariant Graph Neural Network Model for Predicting the Structures of Protein Mutants with Single-Site Mutations

We designed and developed a deep learning model (PreMut) based on an Equivariant Graph Neural Network (EGNN) [32] to predict the tertiary structure of a protein mutant with a single-site mutation (Figure 1). The input is a graph representation of the initial input structure of a mutant constructed from the known structure of its wild-type counterpart, where each node denotes an atom and an edge connects a node with each of its K nearest neighbors (K = 30). PreMut transforms the node features that are initialized as types of atoms through the EGNN in a fashion that is equivariant to the rotation and translation of the positions of nodes (atoms) in the input structure. The transformed node features are used to predict the structural changes caused by the mutation, which are added to the input structure via a residual connection [33] to generate a predicted tertiary structure for the mutant.

**Fig. 1:**
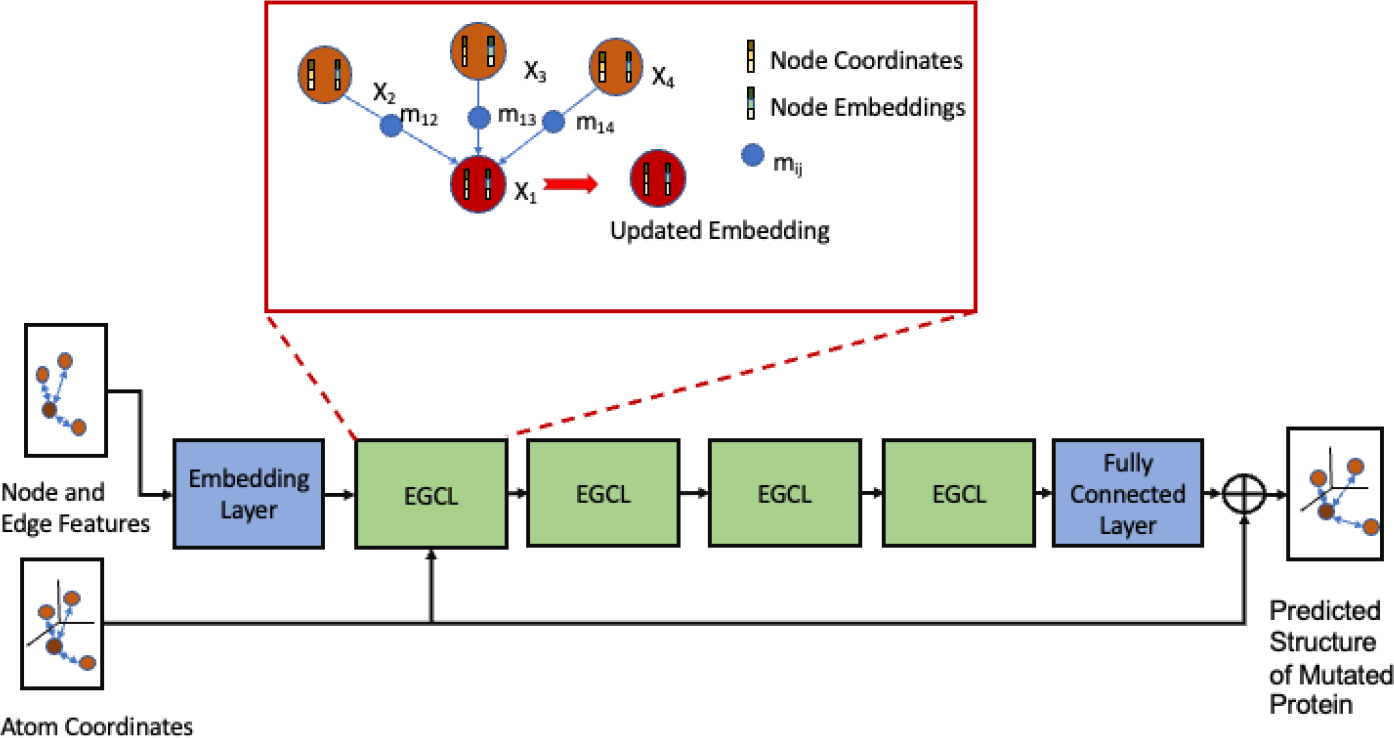
The overall architecture of PreMut for predicting the structure of a mutated protein with a single-site mutation. The input is a graph representation of the initial structure of a protein mutant constructed (essentially copied) from its wild-type counterpart. The node features of the input graph are first processed by an embedding layer to generate node embeddings. These node embeddings along with node coordinates and edge features (i.e., the distances between connected nodes) are used as input for four equivariant graph convolutional layers (EGCL) to generate the hidden node features (embeddings) that are translation- and rotation-equivariant to the coordinates as well as the updated coordinates of the nodes. The hidden features of each node generated in the last EGCL layer are used as input for a fully connected layer to predict the structural change of the position of the node, which is added to the initial input coordinates of the node via a residual connection to generate the x, y, and z coordinates of each node of the mutated protein, resulting in a predicted structure. The transformation of the hidden node features of the graph via an EGCL is briefly illustrated in the red box, where a message *m*_*ij*_ is produced through an edge operation on each node *i* and each of its neighboring nodes (*j*) from their coordinates and embeddings in an rotation- and translation-equivariant way. The messages from all its neighboring nodes and its hidden features are used to generate the new features (embeddings) for the node. More details about the feature transformation are described in the Materials and Methods Section.

After PreMut was trained and validated on the training and validation data, we benchmarked it against a baseline method (AlphaFold) on two independent test datasets (MutData2022 test and MutData2023 test) whose mutant proteins have less than 30% sequence identity with any protein mutant in the training and validation data. MutData2022 test has 75 protein mutants that are clustered in 22 clusters according to the 30% sequence identity threshold. The structures of the mutants in MutData2022 test were released in the PDB by 19th September 2022. MutData2023 test has 59 mutants that are clustered into 10 clusters according to the 30% sequence identity threshold. The structures of the mutants in MutData2023 test were released in the PDB in 2023 after the training data for PreMut were curated. For each method, we measured its average performance (e.g., TM-score [35] of predicted structure) over all the targets in a dataset (e.g., per-target average TM-score) as well as all the clusters (e.g., per-cluster average TM-score). The former is a simple average of the scores of all the targets in the data without considering the size of clusters so that a large cluster with a lot of similar proteins has a high weight in the calculation of the average. The latter first computes the average score of the targets in each cluster in a dataset and then calculates the average of the average scores of all the clusters in the dataset as the result. Therefore, the average per-cluster score weighs the clusters equally regardless of their sizes.

### 2.2 Comparing PreMut with AlphaFold2 in terms of the quality of predicted backbone structures

The structures for the mutated proteins in each test dataset (Mut-Data2022 test or MutData2023 test) predicted by each method (PreMut and AlphaFold) were evaluated against their experimental structures in terms of the two widely used metrics (GDT-HA [36] and TM-score [35]). TM-score or GDT-TS score [36] assesses the topological similarity between two protein structures, while GDT-HA is a high-resolution variant of GDT-TS that is more sensitive to the minor difference between similar protein structures than GDT-TS score or TM-score. In this work, the two complementary metrics GDT-HA and TM-score were used to evaluate the performance of the two methods. It is worth noting that both metrics only assess the quality of the backbone atoms of a protein structure without considering side-chain atoms.

Table 1 reports the per-cluster average GDT-HA scores and TM-scores of PreMut and AlphaFold on MutData2022 test and MutData2023 test. On MutData2022 test, PreMut outperforms AlphaFold in terms of both average per-cluster TM-Score and GDT-HA score, and on MutData2023 test, PreMut outperforms Alphafold. For instance, on MutData2022 test, the per-cluster average GDT-HA score of PreMut is 0.9272, 8.2% higher than 0.8568 of AlphaFold. On MutData2023 test, the per-cluster average GDT-HA score of PreMut is 0.8128, 6.8% higher than 0.7611 of AlphaFold. PreMut also outperforms AlphaFold in terms of TM-score, even though the difference is smaller than in terms of GDT-HA score because the TM-score is less sensitive to small variations between similar structures. The results also show that refining the structures predicted by PreMut using ATOMRefine [30] (i.e., PreMut (Refined)) yields almost the same scores as PreMut, indicating that the refinement maintains the good quality of the backbone structures predicted by PreMut.

**Table 1:** The per-cluster average GDT-HA and TM-score of PreMut, AlphaFold on MutData2022 test and MutData2023 test. Bold font denotes the highest score.

|  | MutData2022_test |  | MutData2023_test |  |
| --- | --- | --- | --- | --- |
| Model | GDT-HA Score | TM-score | GDT-HA Score | TM-score |
| AlphaFold | 0.8568 | 0.9508 | 0.7611 | 0.9489 |
| PreMut | **0.9272** | **0.9767** | **0.8128** | **0.9589** |
| PreMut (Refined) | 0.9268 | 0.9766 | **0.8128** | **0.9589** |

Table 2 reports the per-target average GDT-HA score and TM-score of the three methods on the two datasets. Similarly, PreMut or PreMut (Refined) performs better than AlphaFold on the two datasets in terms of both the per-target average GDT-HA score and the per-target average TM-score. The t-test on the difference between the per-target average GDT-HA scores of PreMut/PreMut(Refined) and AlphaFold indicates that PreMut or PreMut(Refined) significantly outperforms AlphaFold according to the p-value threshold of 0.05 in most situations (e.g., the p-value of the difference between PreMut and AlphaFold = 0.0000008 on GDT-HA score for per target average and p-value = 0.0049884 for per cluster average on MutData2022 test dataset).

**Table 2:** The per-target average GDT-HA score and TM-score of PreMut, AlphaFold on MutData2022 test and MutData2023 test. Bold font denotes the highest score.

|  | MutData2022_test |  | MutData2023_test |  |
| --- | --- | --- | --- | --- |
| Model | GDT-HA Score | TM-score | GDT-HA Score | TM-score |
| AlphaFold | 0.8854 | 0.9688 | 0.8151 | 0.9625 |
| PreMut | **0.9411** | **0.9857** | **0.9396** | **0.9862** |
| PreMut (Refined) | 0.9411 | 0.9856 | **0.9396** | **0.9862** |

Figure 2 shows the polygonal histograms of the GDT-HA scores of the two methods on MutData2022 test and MutData2023 test. The histograms show the GDT-HA scores of PreMut are more concentrated in the high-score range [0.8, 1] than AlphaFold on the two datasets. On MutData2023 test, a significant portion of the structures predicted by AlphaFold has a medium GDT-HA score between 0.6 and 0.8.

**Fig. 2:**
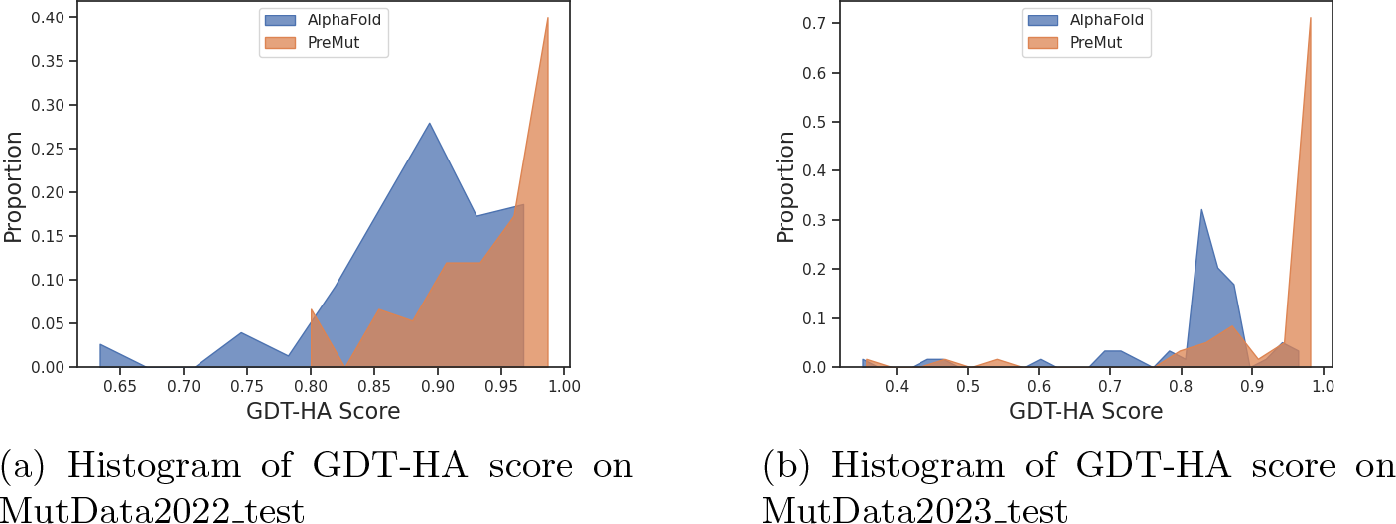
The polygonal histogram plots of GDT-HA scores of PreMut and AlphaFold on MutData2022 test and MutData2023 test.

The scatter plots in Figure 3 plot the GDT-HA score of the structure of each mutant predicted by PreMut against that of AlphaFold on MutData2022 test and MutData2023 test, respectively. The plots show that PreMut has a higher score on more mutants than AlphaFold on each dataset. For instance, on Mut-Data2022 test, PreMut has a higher score on 58 targets than AlphaFold, while AlphaFold only has a higher score on 17 targets; on MutData2023 test, Pre-Mut has a higher score on 52 targets than AlphaFold, while AlphaFold only has a higher score on 7 targets.

**Fig. 3:**
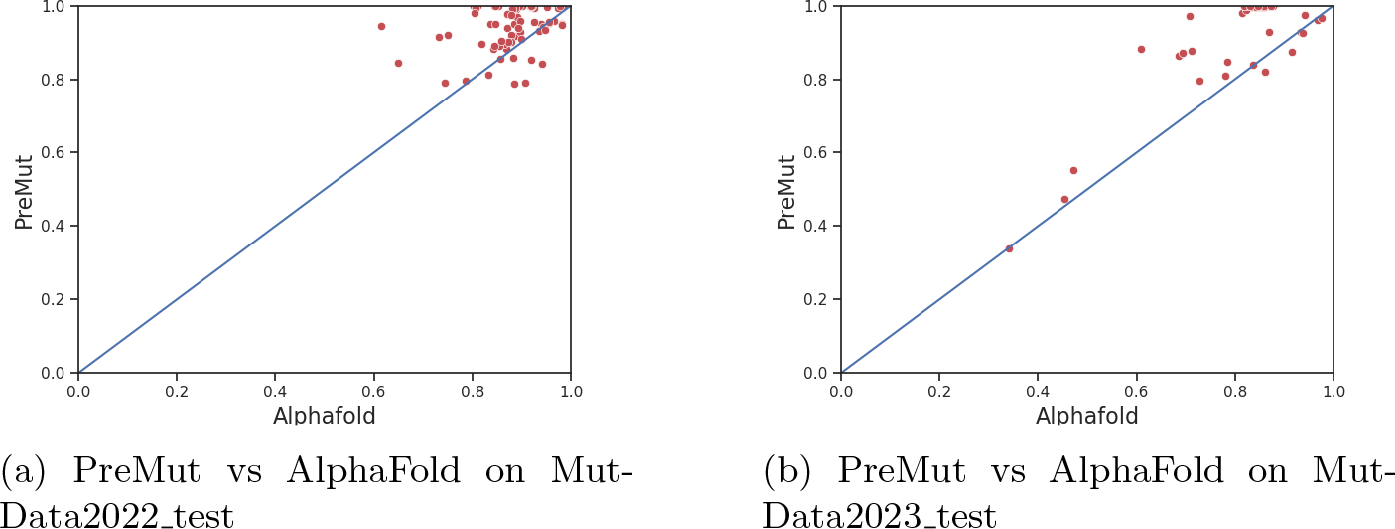
The scatter plots of the GDT-HA scores of the mutant structures predicted by PreMut against those predicted by AlphaFold on MutData2022 test and MutData2023 test, respectively. A dot on the 45-degree diagonal line indicates the structures predicted by two methods for a target have the (almost) same score, while a dot above indicates the line PreMut outperforms the other method on a target. (a) The scatter plot of PreMut against AlphaFold, where PreMut performs better than AlphaFold on 58 out of 75 targets in MutData2022 test. (b) The scatter plot of PreMut against AlphaFold, where PreMut performs better than AlphaFold on 52 of 59 targets in Mut-Data2023 test.

### 2.3 Comparing PreMut with AlphaFold in terms of quality of side-chains of predicted structures

We used the Superposition-based Protein Embedded Carbon-Alpha Side-chain (SPECS) score [37] to evaluate the side-chains of the structures of the mutants predicted by PreMut and AlphaFold. SPECS measures the similarity between the side-chains of a predicted structure and those of the native (experimental) structure. It utilizes the united-residue-representation [38] to calculate global carbon-alpha positioning-based distance, side-chain distance, and side-chain orientation and combine them to measure the side-chain similarity. The per-target average SPECS scores and per-cluster average SPECS scores of the methods are reported in Table 3. The results show that PreMut or PreMut (Refined) performs better than AlphaFold. For instance, on MutData2022 test, the per-target average SPECS score of PreMut (Refined) is 0.7024, higher than 0.6328 of AlphaFold; on MutData2023 test, the per-target average SPECS score of PreMut is 0.7011, higher than 0.5896 of AlphaFold.

**Table 3:** The average SPECS scores of predicted mutant structures on MutData2022 test and MutData2023 test. Bold font denotes the best result.

| Method | Per-Target Average |  | Per-Cluster Average |  |
| --- | --- | --- | --- | --- |
|  | MutData2022_test | MutData2023_test | MutData2022_test | MutData2023_test |
| AlphaFold | 0.6329 | 0.5896 | 0.6143 | 0.5514 |
| PreMut | 0.6727 | 0.6745 | 0.6629 | 0.5755 |
| PreMut (Refined) | **0.7024** | **0.7011** | **0.6927** | **0.6029** |

Moreover, the SPECS score of PreMut (Refined) is higher than PreMut on both datasets in terms of both the per-target average and the per-cluster average indicating that using AtomRefine to refine the structures predicted by PreMut improves the quality of the side chains, which is different from their almost same performance in terms of the quality of the backbone structures (Table 1).

Figure 4 plots the SPECS score of PreMut (Refined) against AlphaFold on MutData2022 test and MutData2023 test, respectively. The plots show that PreMut (Refined) has higher SPECS scores for a majority of the mutants than AlphaFold on the two datasets.

**Fig. 4:**
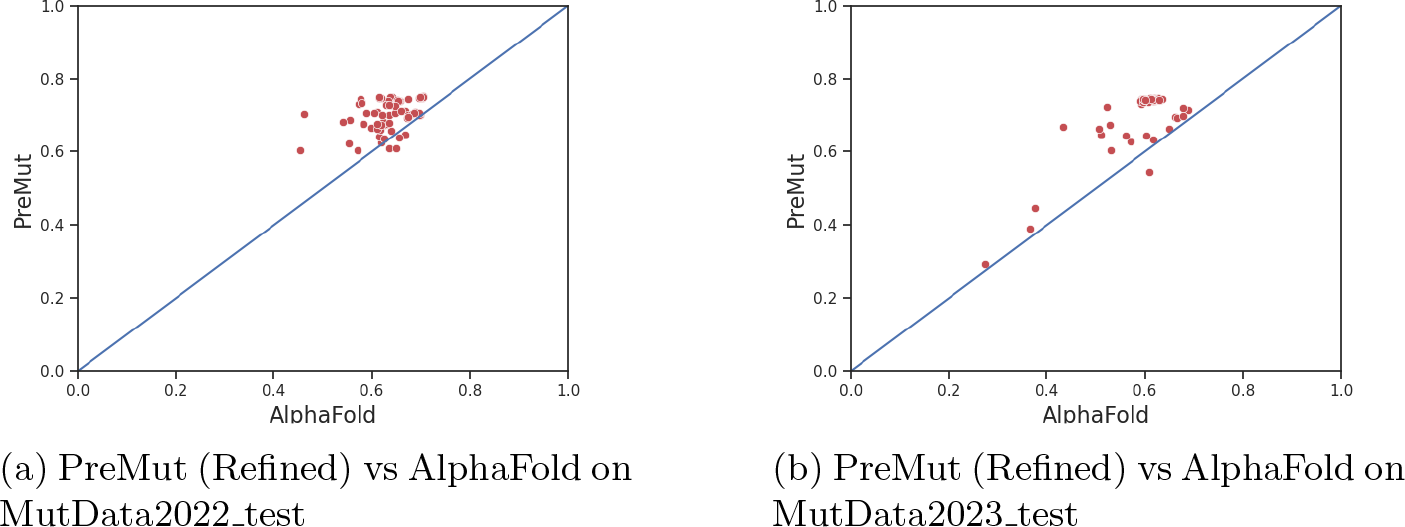
The scatter plots of the SPECS scores of PreMut (Refined) against AlphaFold on MutData2022 test and MutData2023 test. A dot on the 45-degree diagonal line indicates the structures predicted by two methods for a target have the (almost) same score, while a dot above the line indicates PreMut (Refined) outperforms AlphaFold on a target. (a) The plot of PreMut (Refined) against AlphaFold where PreMut (Refined) performs better on 71 out of 75 targets in MutData2022 test. (b) The plot of PreMut (Refined) against AlphaFold where PreMut (Refined) performs better on 58 out of 59 targets in MutData2023 test.

### 2.4 An Ablation Study of Node Features

We tested two different kinds of features describing each atom (node) in the graph representing the structure of a protein mutant. One is the one-hot encoding of 37 possible types of the atom observed in the PDB. Another one is to combine the one-hot encoding of atom types and the encoding of the type of the residue that the atom belongs to. The residue type is a vector consisting of 21 numbers corresponding to the 20 standard amino acids and an unknown amino acid type. For a non-mutated position, all numbers are set to 0 except the one denoting the actual amino acid type of the position is set to 1. For the mutated position, the number denoting the mutated residue is set to 1, the number denoting the original amino acid is set to -1, and all other numbers are set to 0.

Table 4 and Table 5 report the per-cluster and per-cluster average TM-scores and GDT-HA scores of the two kinds of node features. They results show that they have very similar performance, indicating that a simple representation of the type of each atom is sufficient for PreMut to predict the structural changes of proteins caused by single-site mutations. Therefore, only the atom type feature is used in the final version of PreMut.

**Table 4:** The per-cluster average TM-scores and GDT-HA scores of two kinds of features on MutData2022 and MutData2023. Bold font denotes the highest score.

|  | MutData2022_test |  | MutData2023_test |  |
| --- | --- | --- | --- | --- |
| Features | TM-score | GDT-HA Score | TM-score | GDT-HA Score |
| Atom Type Only | **0.9767** | **0.9272** | 0.9589 | 0.8128 |
| Atom Type + Residue Type | 0.9765 | 0.9270 | **0.9610** | **0.8199** |

**Table 5:** The per-target average TM-scores and GDT-HA scores of two kinds of features on MutData2022 test and MutData2023 test. Bold font denotes the highest score.

|  | MutData2022_test |  | MutData2023_test |  |
| --- | --- | --- | --- | --- |
| Model | TM-score | GDT-HA Score | TM-score | GDT-HA Score |
| Atom Type Only | **0.9857** | **0.9411** | 0.9862 | 0.9396 |
| Atom Type + Residue Type | 0.9856 | 0.9409 | **0.9865** | **0.9408** |

### 2.5 A Case Study

We investigated the structure prediction for a specific protein mutant, i.e., chain A of protein 7MGR in the PDB [39], which is one residue different from the chain A of a wild protein 8B0S [40] in the PDB, to illustrate where PreMut may perform better than AlphaFold. They are the main protease of severe acute respiratory syndrome coronavirus 2 (SARS-CoV-2). At position 145, residue C in the wild protein is replaced by residue A in the mutant (i.e., C145A). The GDT-HA score of the structure predicted by PreMut (Refined) for the mutant is 0.8829, 45.21% higher than 0.6080 of AlphaFold. The TM-score of PreMut (Refined) is 0.9911, 21.58% higher than 0.8145 of AlphaFold. The SPECS score of PreMut (Refined) is 0.6699, 54.21% higher than that of 0.4344 of AlphaFold.

Figure 5a and Figure 5b show the superimposition between the structure predicted by AlphaFold/PreMut (Refined) and the experimental structure of the mutant. The structures are visualized by Chimera [41]. It is shown that the overall structure of the structure predicted by PreMut (Refined) aligns well with the experimental structure, while there is some obvious difference between that by AlphaFold and the experimental structure, particularly in the rightmost small alpha helical domain.

**Fig. 5:**
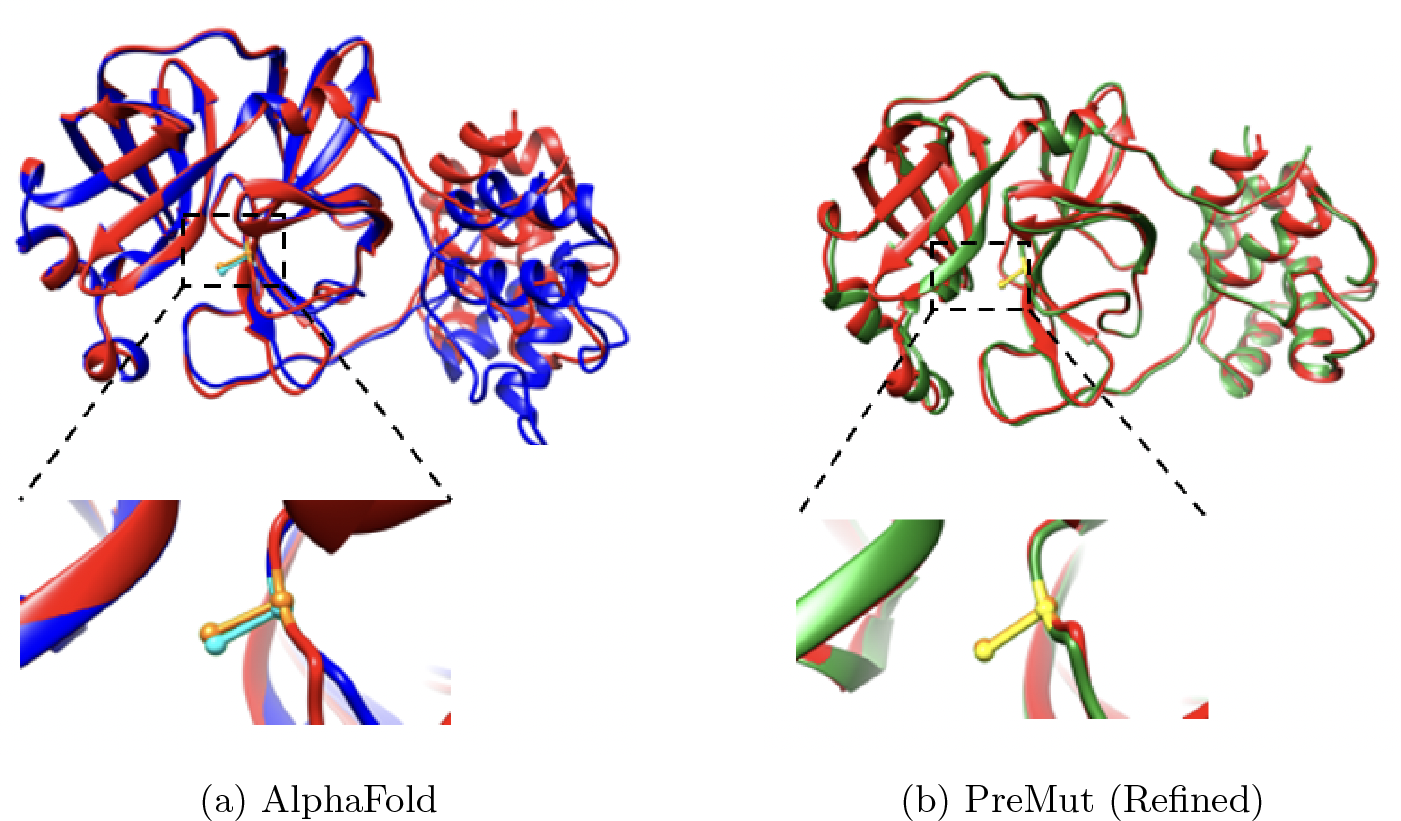
The superimposition of the predicted structure for a mutant (the chain A of 7MGR with a single-site mutation C145A) and its experimental (native) structure. (a) AlphaFold predicted structure in blue superimposed with the experimental structure in red. The side-chain atoms of the mutated residue are shown as ball and stick in the zoom-in view (orange stick: side-chain atoms in the native structure; cyan sticks: side-chain atoms in the AlphaFold prediction). (b) PreMut (Refined) predicted structure in green superimposed with the experimental structure in red. In the zoom-in view, the orange stick denotes the side-chain atoms in the native structure and the yellow stick the side-chain atoms in the PreMut (Refined) prediction.

Zooming in on the region around the mutated residue, the backbone at the mutated position in the structure predicted by PreMut (Refined) aligns more tightly with the nature structure (Figure 5b) than in the structure predicted by AlphaFold (Figure 5a). Moreover, the residues near the mutated position in the structure predicted by PreMut (Refined) align well with those in the experimental structure, while some residues near the position in the structure predicted by AlphaFold deviate substantially from those in the experimental structure.

## 3 Conclusion and Future Work

Predicting the structures of proteins with single-site mutations is important for studying protein function, understanding the genetic underpinnings of phenotypes, and advancing protein engineering. However, existing mutation prediction methods can only predict the stability or function effects of mutations without directly predicting the tertiary structures of mutated protein structures. The general protein structure prediction methods such as AlphaFold are not sensitive enough to predict the structural changes induced by a single amino acid mutation. To address this gap, we have introduced an equivariant graph neural network method (PreMut) to predict the protein tertiary structural change caused by single-site mutations with respect to the structure of the wild protein and consequently the structure of the mutated protein. PreMut can more accurately predict both the backbone and side-chain structures of mutated proteins than the state-of-the-art deep learning protein structure prediction tool - AlphaFold. Therefore, our experiment demonstrates that the protein tertiary structural change of many proteins induced by mutations can be rather accurately predicted by deep learning methods. However, one limitation of PreMut is that it still fails to predict the structure of some mutants. As there is still some gap between the predicted structures of mutants and their native structures, we expect more sophisticated deep learning methods will be developed in this direction. Another limitation is that PreMut can only predict the structures of mutants with single-site mutations. In the future, we plan to extend it to predict the structures of mutants with an arbitrary number of mutations.

## 4 Materials and Methods

### 4.1 Protein Mutation Data Curation

We downloaded all the protein structures available in the Protein Data Bank (PDB) as of 19th September 2022. The protein structures with mutations indicated by the keyword ‘ENGINEERED MUTATION’ in the lines starting with the word ‘SEQADV’ in their PDB files were retained. The PDB files of the mutants contain the mutant information including the positions of mutations, original residues, and substitute residues. We only considered mutants consisting of single-site mutations.

In order to find the wild protein counterparts for the mutants collected above, the sequence of the wild protein for each mutant was generated by replacing the mutated residue in the sequence of the mutant with the original residue. The sequence of the wild protein was searched against the sequences of all the proteins in the PDB by mmseqs[42] to identify the proteins that have 100 percent identity with the wild protein sequence. If such a protein was found, it was paired with the mutant to form a wild protein-mutant pair. In this process, we found that there can be a many-to-many pairing relationship between wild proteins and mutants, i.e., a mutant may have multiple wild proteins in the PDB that have the same sequence and a wild protein may also correspond to multiple mutants. In total, 36,458 pairs of wild and mutated proteins were collected, which involved 1,094 unique mutants with distinct PDB codes.

To reduce the redundancy of the pairs, we used mmseqs to cluster the sequences of mutants according to the 30% sequence identity threshold, i.e., the pairs whose mutant protein sequence has at least 30% sequence are grouped into one cluster. In the end, the 36,458 pairs of wild and mutant proteins were grouped into 225 clusters. This dataset is called MutData2022.

MutData2022 was randomly split into the training/validation data and the test data according to the 90% and 10% ratio, resulting in 203 clusters for training/validation and 22 clusters for testing (called MutData2022 test). The training/validation data was then randomly split into training data (182 clusters) and validation data (21 clusters) according to the 90% and 10% ratio. The training data was used to optimize the parameters of the deep learning models, while the validation data was used to select the best model for the final test. It is worth noting that during the training process, the wild protein structure and the mutant structure of each pair in the training dataset were superimposed by TMAlign [43] to make them in the same coordinate system so that PreMut can be trained to focus on learning to predict the difference between the input structure of a mutant constructed from the wild-type protein structure and the true (experimental) structure of the mutant.

To create another independent test dataset, we applied the same strategy above to the proteins in the PDB released by May 2023, excluding all the proteins present in MutData2022. The dataset is called MutData2023 test. This dataset contains mutants released after the curation of MutData2022. To remove the redundancy with MutData2022, we utilized the mmseqs-based filtering to make sure that every mutant in MutData2023 test did not have more than 30% of sequence identity with any mutant in MutData2022. The wild and mutated protein pairs in MutData2023 test were clustered by mmseqs into 10 clusters consisting of 59 pairs in total according to the 30% sequence identity threshold on the sequences of the mutants.

### 4.2 Graph Representation of Protein Structures, Input Features, and Labels

As in [30], a graph is used to represent all the atoms of an initial input structure for a mutant, in which a node denotes an atom and an edge connects a node with its neighbors. The closest 30 atoms for each atom (node) were selected by the K-Nearest Neighbour algorithm[44] as its neighbors, resulting in *N ×* 30 edges (*N* : the number of nodes). Each edge has one feature - the Euclidean distance between the two nodes of the edge.

The initial input structure for a mutant is constructed from the structure of its corresponding wild-type protein as follows. The wild protein structure is first centralized by subtracting the x, y, or z coordinates of each atom by the mean of x, y, or z coordinates of all its atoms. In the training phase, the centralized coordinates of the wild protein structure were superimposed with the structure of the mutant using TMAlign to make them in the same coordinate system to facilitate the deep learning method to predict the structural change from the wild-type protein structure to the structure of the mutant. But this superimposition step is not needed in the prediction phase at all. The coordinates of the atoms of all the residues except the mutated one of the initial input structure for a mutant are copied from the structure of the wild-type protein. The atoms of (e.g., mostly the backbone atoms C, Ca, N, O) of the mutated residue of the initial input structure are also copied from their counterparts in the wild protein structure if they exist. The coordinates of the atoms (e.g., side-chain atoms) of the mutated residue of the initial input structure that they do not have counterparts in the wild protein structure are randomly initialized as a position within 1.54 Angstrom [45] of its Ca atom.

The type of an atom is used as the node feature invariant to the rotation and translation of protein structures. All 37 types of atoms occurring in the PDB structure files are considered. The one hot encoding of the atom types of the nodes is stored in a matrix with a shape of *N ×* 37, where *N* is the number of atoms.

The x, y, and z coordinates of the atoms of an initial input structure are also used as input for the nodes, which are stored in a matrix of shape *N ×* 3. However, they change with the rotation and translation of input protein structures and therefore are supplied as an input separate from the rotation- and translation-invariant node features such as the atom type features.

The true (experimental) structures of the mutants are used as the targets (i.e., labels) to train the deep learning method to predict the structures of the mutants from their input structures.

### 4.3 Design and Training of the Equivariant Graph Neural Network and Structure Refinement

#### 4.3.1 Embedding Layer

As shown in Figure 1, the first layer of PreMut is an embedding layer - a fully connected neural network layer, which converts the node features of the input graph representing the initial structure of a mutant with a shape of *N ×* 37 (*N* : number of nodes) to the node embeddings with a shape of *N ×* 32.

#### 4.3.2 Equivariant Graph Convolutional Layer (EGCL)

PreMut has four EGCL layers to update the embeddings of the nodes in a protein structure graph and their coordinates (Figure 1). The inputs to an EGCL layer *l* are the node embeddings *h*^*l*^, node coordinates *x*^*l*^, and edge features *e*^*l*^ from the previous layer. It generates the updated node embeddings and coordinates *h*^*l*+1^, *x*^*l*+1^ = *EGCL*(*h*^*l*^, *x*^*l*^, *e*^*l*^) using the four equations below [32].

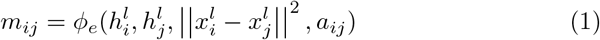

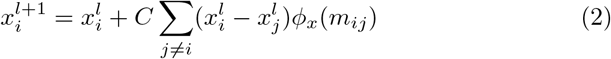

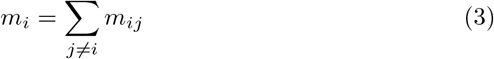

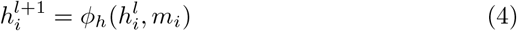

Equation 1 applies an edge operation *ϕ*_*e*_ (a neural network function) to the current embedding of each node 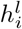, the embedding of one of its neighboring nodes 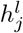, the squared distance between two nodes 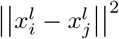 and the features of the edge connecting the two nodes *a*_*ij*_ to generate a message from node *j* to node *i*: *m*_*ij*_. Here, the *ϕ*_*e*_ is a multilayer perceptron (MLP) that has one hidden layer and an output layer with the activation function of the Sigmoid Linear Unit (SiLU) [46]. The output of the MLP is used by an attention layer (a feedforward neural network with a Sigmoid function) [47] to produce the attention weights, which are multiplied with the MLP’s output to generate the final output of the edge operation. Equation 2 generates the new coordinates for node *i* from its current coordinates and the messages from its neighboring nodes. *ϕ*_*x*_ is an MLP consisting of one hidden layer and an output layer with the activation function of SiLU. Here, *C* is equal to 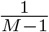, where *M* is the number of neighboring nodes. Equation 3 calculates the total message to node *i* from all its neighbors. Finally, Equation 4 computes the updated embedding for node *i* from its current embedding and the total message from its neighbors. *ϕ*_*h*_ is an MLP consisting of a single hidden layer and an output layer with the activation function of SiLU. The dimension of the updated embedding for node *i* in each EGCL layer is 32.

#### 4.3.3 Predicting the Structure of a Protein Mutant

The embedding of each node in the last ECGL layer is used as input for a fully connected neural network layer with a linear activation function in Figure 1 to predict the change of the x, y, z coordinates of the node upon a single-site mutation, i.e., the difference between the true structure of a protein mutant and the structure of its wild-type counterpart (i.e., the input structure), which is added to the initial input structure via a linear residual connection to generate the final predicted structure of the mutant whose x, y, z coordinates are stored in a tensor with a shape of *N ×* 3.

#### 4.3.4 Training

We treat the prediction of the structure of a protein mutant as a regression problem. In the training process, the mean squared difference between the coordinates of the predicted structure of a mutant and those of its true (experimental) structure was calculated as the loss.

PreMut was trained on the training data for 100 epochs using the Adam optimizer [48] with a learning rate of 0.0001 to minimize the loss. During each epoch, the structural similarity between the predicted structure and the true structure (i.e. TM-score) on the validation dataset was calculated, which was used to select the final trained model for the blind test.

#### 4.3.5 Refinement of Predicted Structures

To remove possible atom-atom clashes and violations of bond lengths and angles in a predicted protein structure, we apply a protein structure refinement tool ATOMRefine [30] to refine it to generate a refined protein structure. This optional refinement step can improve the quality of side-chain atoms in the predicted structure while maintaining the similar quality of backbone atoms (see the Results Section).

### 4.4 AlphaFold for Mutant Structure Prediction

We used AlphaFold as a baseline method for protein mutant structure prediction to benchmark PreMut. We used its online version Colabfold [49] to predict the structure of a mutated protein. We used Colabfold to take a sequence of a protein mutant as the sequence input and the known structure of its wild-type protein as the template input to predict the structures for the mutant. The no. 1 ranked structure predicted by Colabfold was used as the predicted structure for the mutant in the evaluation.

## Declarations

### Funding

The work was partly supported by the National Science Foundation (grant #: DBI2308699) and the National Institutes of Health (grant #: R01GM093123 and R01GM146340).

### Conflict of interest

The authors declare there is no conflict of interest.

### Consent for publication

All the authors are consent for publication.

### Availability of data and materials

The training and test data are available at https://zenodo.org/record/8401256.

### Code availability

The source code of PreMut is available at https://github.com/jianlin-cheng/PreMut

## Acknowledgements

We would like to thank Dr. Jack Tanner for valuable discussions.

## Authors’ contributions

JC conceived and supervised the project. JC and SM designed the method and the experiment. SM implemented, trained, and tested the method, collected the data, and analyzed the data. AM made some contributions to data processing and training. SM and JC wrote the manuscript.

## Notes

https://zenodo.org/record/8401256

https://github.com/jianlin-cheng/PreMut

### Competing Interest Statement

The authors have declared no competing interest.

### Summary of Updates

Fixed typos and URL links.

